# Gene2Function: An integrated online resource for gene function discovery

**DOI:** 10.1101/133975

**Authors:** Yanhui Hu, Aram Comjean, Stephanie E. Mohr, the FlyBase Consortium, Norbert Perrimon, Julie Agapite, Kris Broll, Madeline Crosby, Gilberto Dos Santos, David Emmert, Kathleen Falls, Susan Russo Gelbart, Sian Gramates, Beverley Matthews, Norbert Perrimon, Carol Sutherland, Chris Tabone, Pinglei Zhou, Mark Zytkovicz, Giulia Antonazzo, Helen Attrill, Nicholas Brown, Silvie Fexova, Tamsin Jones, Aoife Larkin, Steven Marigold, Gillian Milburn, Alix Rey, Jose-Maria Urbano, Brian Czoch, Josh Goodman, Gary Grumbling, Thomas Kaufman, Victor Strelets, James Thurmond, Phill Baker, Richard Cripps, Margaret Werner-Washburne

**Affiliations:** Drosophila RNAi Screening Center, Dept. of Genetics, Harvard Medical School, Boston, MA; The FlyBase Consortium; Howard Hughes Medical Institute; Harvard University; University of Cambridge, UK; Indiana University; University of New Mexico

## Abstract

One of the most powerful ways to develop hypotheses regarding biological functions of conserved genes in a given species, such as in humans, is to first look at what is known about function in another species. Model organism databases (MODs) and other resources are rich with functional information but difficult to mine. Gene2Function (G2F) addresses a broad need by integrating information about conserved genes in a single online resource.

## Main text

The availability of full-genome sequence has uncovered a striking level of conservation among genes from single-celled organisms such as yeast; invertebrates such as flies or nematode worms; and vertebrates such as fish, mice, and humans. This conservation is not limited to amino acid identity or structure, or RNA sequence. Indeed, gene conservation often extends to conservation of biochemical function (e.g. common enzymatic functions); cellular function (e.g. specific role in intracellular signal transduction); and function at the organ, tissue, and whole-organism levels (e.g. control of organ formation, tissue homeostasis, or behavior).

Researchers applying small-and large-scale approaches in any common model organism often come across genes that are poorly characterized in their species of interest. A common and powerful way to develop an hypothesis regarding the function of a gene poorly characterized in one species—or newly implicated in some processes in that species—is to ask if the gene is conserved and if so, find out what is known about the functions of its orthologs in other species. This commonly applied approach gains emphasis when the poorly characterized gene is implicated in a human disease; in many cases, what we know about human gene function is largely based on what was first uncovered for orthologs in other species.

Despite the importance and broad application of this approach among biologists and biomedical researchers, there are barriers to applying the approach to its fullest. First, ortholog mapping is not straightforward. Over the years, many approaches and algorithms have been applied to mapping of orthologs. The results do not always agree and at a practical level, the use of different genome annotation versions, as well as different gene or protein identifiers, can make it difficult to identify or have confidence in an ortholog relationship. Second, even after one or more orthologs in common model species have been identified, it is not easy to quickly assess in which species the orthologs have been studied and determine what functional information was gained. Model organism databases (MODs) and human gene databases provide relevant, expertly curated information. Although InterMine ^1^ provides a mechanism for batch search of standardized information and NCBI Gene provides information about individual genes in a standardized format, it remains a challenge to navigate, access, and integrate information about all of the orthologs of a given gene in well-studied organisms. As a result, useful information can be missed, contributing to inefficiency and needless delay in reaching the goal of functional annotation of genes, including human disease-relevant genes.

Clearly, there is a need for an integrated resource that facilitates the identification of orthologs and mining of information regarding ortholog function, in particular in common genetic model organisms supported by MODs. Previously, we developed approaches for integration of various types of gene-or protein-related information, including ortholog predictions (DIOPT; ^2^), disease-gene mapping based on various sources (DIOPT-DIST; ^2^), and transcriptomics data (DGET; ^3^). Importantly, these can serve as individual components of a more comprehensive, integrated resource. Indeed, our DIOPT approach to identification of high-confidence ortholog predictions is now used in other contexts, including at FlyBase ^4^ and at MARRVEL for mining information starting with human gene variant information (^5^; www.marrvel.org).

To address the broad need for an integrated resource, we developed www.gene2function.org (G2F), an online resource that maps orthologs among human genes and common genetic model species supported by MODs, and displays summary information for each ortholog. G2F makes it easy to survey the wealth of information available for orthologs and easy to navigate from one species to another, and connects users to detailed reports and information at individual MODs and other sources. The integration approach and set of information sources are outlined in ***Figure 1*** and described in the ***Supplemental Methods***.

**Figure 1:**
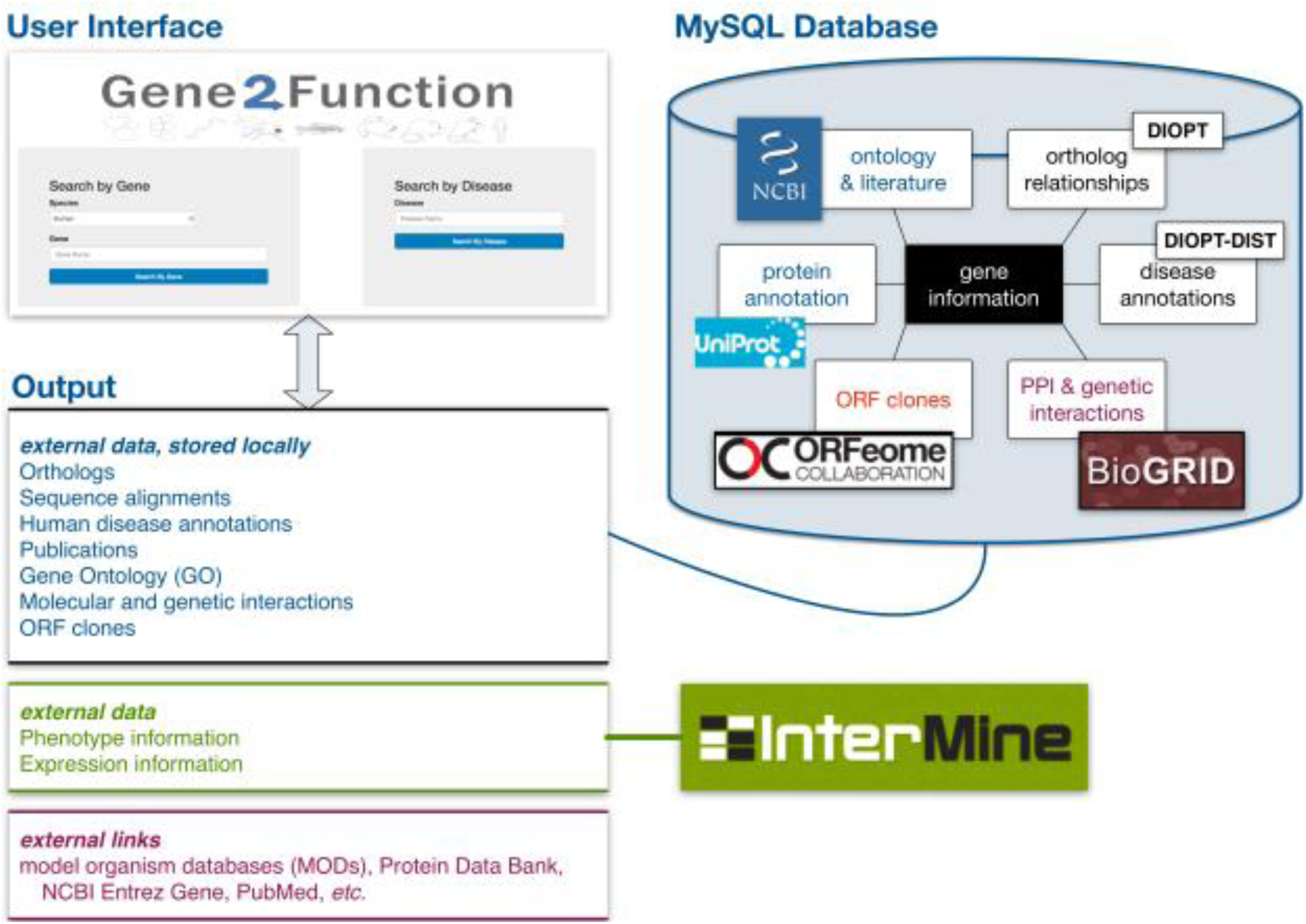
Overview of the Gene2Function (G2F) online resource. For detailed information about the database, logic flow, and information sources, see ***Supplemental Methods***.

To demonstrate the utility of G2F, we focus on two use cases, (1) a search initiated with a single human or common model organism gene of interest, and (2) a search initiated with a single human disease term of interest (***Fig. 2***).

**Figure 2:**
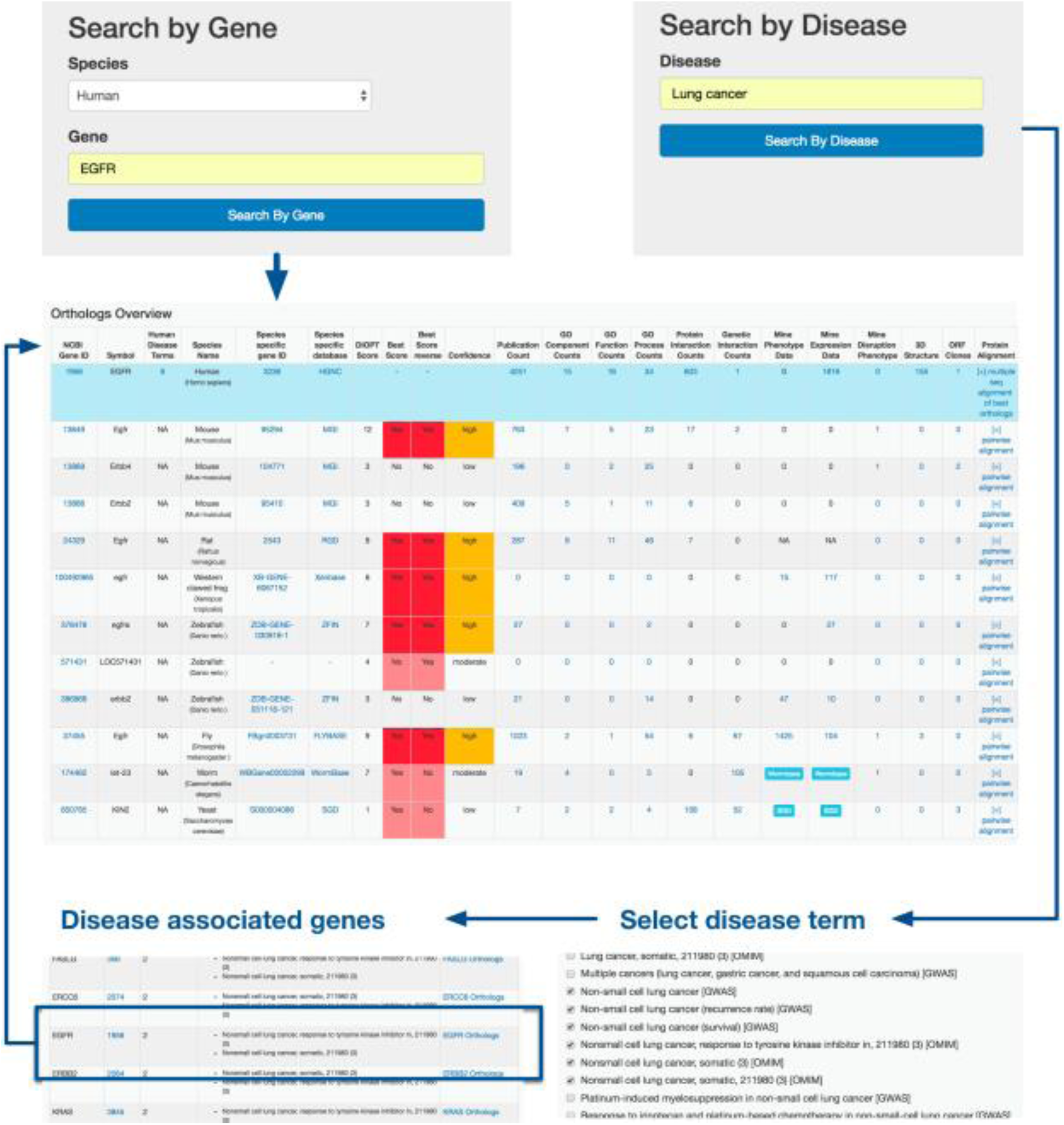
Gene and disease search user interfaces at G2F. The results of a gene search are displayed as a table of orthologs that summarizes and links to functional and other information, a multi-sequence alignment, and pairwise alignments. The results of a disease search first allow for disambiguation of terms, then display a table of genes associated with the term(s). Each disease-associated gene links to the same summary table displayed for a gene search.

A gene search at G2F connects users to ortholog information and an overview of functional information for orthologs. Specifically, starting with a search of a human, mouse, frog, fish, fly, worm, or yeast gene, users reach a summary table of orthologs and information (***Fig. 2***).

Information displayed includes the number of gene ontology (GO) terms assigned based on experimental evidence; the number of publications; and the number of molecular and genetic interactions reported. When available, the table also includes links to expression pattern annotations, phenotype annotations, three dimensional structure information ^6^, and open reading frame (ORF) clones from the ORFeome collaboration consortium ^7-9^ that are available in a public repository ^10^. The summary allows a user to quickly 1) evaluate conservation across major model organisms based on DIOPT score, pairwise alignment of the query protein to another species, and multiple-sequence alignment; 2) assess in what species the query gene has been well-studied based on original publications, annotation, and data; and 3) identify reagents for follow up studies. The summary table also allows a user to view detailed reports and is hyper-linked to more detailed information at original sources, such as data on specific gene pages at MODs.

A disease search at G2F first connects from disease terms to associated human genes, then uses the gene search results table format to display orthologs of the human gene and summary information (***Fig. 2***). After a search with a human disease term, users are first shown a page that helps disambiguate terms, expanding or focusing the search, and also allows users to limit the results to disease-gene relationships curated in the Online Mendelian Inheritance in Man (OMIM) database and/or based on genome-wide association studies (GWAS) from the NHGRI-EBI GWAS Catalog ^11^. Next, users access a table of human genes that match the subset of terms, along with summary information regarding the genes and associated disease terms. On the far right-hand side of the table, users can connect to the same single gene-level report that is described above for a gene search.

Over the past two decades, GWAS have begun to reveal genetic risk factors for many common disorders ^12^. As of Feb 2017, GWAS Catalog ^11^ included 2,385 publications with 10,499 reported genes associated with 1,682 diseases or traits. For some of the human genes, there are no publications or gene ontology annotations. We used G2F to survey information in model organisms for this subset of genes and found many cases where one or more orthologous genes have been studied (***Supplemental Methods***). The results of the ortholog studies appear in some cases to support the disease association and the corresponding model systems could provide a foundation for follow-up studies (***Supplemental Table 1***). The human gene SAMD10, for example, has been shown using the iCOGS custom genotyping array to be one of 23 new prostate cancer susceptibility loci ^13^, but there is no information about this human gene available, aside from sequence and genome location. The results of a G2F search show that the gene is conserved in the mouse, rat, fish, fly, and worm. The mutant phenotypes of the fly ortholog suggest that the gene is involved in compound eye photoreceptor cell differentiation, EGFR signaling, positive regulation of Ras signaling, and ERK signaling, providing starting points for the development of new hypotheses regarding the function of SAMD10. Several uncharacterized human genes associated by GWAS with schizophrenia, namely IGSF9B, NT5DC2, C2orf69, and ASPHD1 ^14^,^15^, are expressed at higher levels in the nervous system than in other tissues in one or more model organisms, suggesting a potential role in the nervous system in these models and supporting the idea that the models might be appropriate for follow-up studies aimed at understanding the human gene function. These examples are extreme in that they represent human genes for which there are no publications describing functional information. For a large number of human genes, limited information is available.

Functional annotations in model systems, as accessed through G2F, can help in the development of new hypotheses regarding the functions of these genes, as well as help researchers choose an appropriate model organism(s) for further study of the conserved gene.

Altogether, G2F provides a highly-integrated resource that facilitates efficient use of existing gene function information by providing a big-picture view of the information landscape and building bridges among different islands of information, including MODs. This approach complements approaches designed for searches starting with long gene lists (e.g. InterMine; ^1^) or based on a phenotype-centered model (e.g. Monarch Initiative; ^16^). The modular nature of the G2F resource makes it possible to easily update the information sources (e.g. replace a module) as well as add new types of information (e.g. an expanded summary of reagents or new types of experimental data).

